# Prediction of Mutations to Control Pathways Enabling Tumour Cell Invasion with the CoLoMoTo Interactive Notebook (Tutorial)

**DOI:** 10.1101/319780

**Authors:** Nicolas Levy, Aurélien Naldi, Céline Hernandez, Gautier Stoll, Denis Thieffry, Andrei Zinovyev, Laurence Calzone, Loïc Paulevé

## Abstract

Boolean and multi-valued logical formalisms are increasingly used to model complex cellular networks. To ease the development and analysis of logical models, a series of software tools have been proposed, often with specific assets. However, combining these tools typically implies a series of cumbersome software installation and model conversion steps. In this respect, the *CoLoMoTo Interactive Notebook* provides a joint distribution of several logical modelling software tools, along with an interactive web Python interface easing the chaining of complementary analyses. In this protocol, we demonstrate the assets of this approach through the analysis of a computational model of biological network. Our computational workflow combines (1) the importation of a GINsim model and its display, (2) its format conversion using the Java library BioLQM, (3) the formal prediction of mutations using the OCaml software Pint, (4) the model checking using the C++ software NuSMV, (5) quantitative stochastic simulations using the C++ software MaBoSS, and (6) the visualisation of results using the Python library matplotlib. Starting with a recent Boolean model of the signalling network controlling tumour cell invasion and migration, our model analysis culminates with the prediction of sets of mutations presumably involved in a metastatic phenotype.

## 1 Introduction

Boolean and multi-valued logical formalisms are increasingly used to model complex cellular networks [5,6,14]. A logical model is usually defined in three steps:

1. The delineation of a regulatory graph, where the vertices (nodes) represent signalling or regulatory components (proteins, genes, miRs, etc.), while the arcs (arrows) represent regulatory interactions between pairs of components. These arcs are labelled by a sign: positive in the case of activation, negative in the case of an inhibition (multiple arcs between two nodes may be considered but are not used here).
2. A discrete variable is associated with each node. In the simplest cases, as hereafter, these variables are Boolean, *i.e*. they can take only two values (0 or 1), denoting the absence/inactivity or the presence/activity of the corresponding components.
3. Finally, a logical rule is associated with each component to specify the combinations enabling its activation. More precisely, this rule combines the different variables corresponding to the regulatory components using the logical negation (noted `|`), conjunction (noted `&`) and disjunction (noted |). For example, the rule associated with the component GF in the model considered below is `!CDH1 & (GF | CDH2)`, which reads as ”the component GF will be activated in the absence of CDH1 and in the presence of CDH2 or GF itself’. In other words, CDH2 is required transiently for GF activation, in the absence of CDH1.

To support the development and analysis of logical models, a series of software tools have been proposed, often with specific assets [7,9,12,13].

The *CoLoMoTo Interactive Notebook*^1^ [11] relies on Docker^2^ and Jupyter^3^ technologies to assist on editing and sharing reproducible analysis workflows for logical models. In addition to the distribution of a set of software tools to define and analyse Boolean and multi-valued networks, a Python interface for each of the integrated tools is provided, greatly easing the execution and chaining of complementary analyses.

This protocol describes in details the usage of the CoLoMoTo Interactive Notebook to provide a reproducible analysis of a recently published model of the signalling network controlling tumour cell invasion and migration. More specifically, we combine different tools (Table 1) to compute the model stable states, perform stochastic simulations, compute (sets of) mutations controlling the reachability of specific stable states, and evaluate their efficiency.

**Table 1:** List of software tools used in this notebook

| Tool | Website | Role in this notebook |
| --- | --- | --- |
| GINsim | ginsim.org | Model input and display, conversion to bioLQM and NuSMV |
| bioLQM | colomoto.org/biolqm | Fixpoint computation, conversion to MaBoSS and Pint |
| MaBoSS | maboss.curie.fr | Stochastic simulations, assess impact of mutations on propensity of reaching phenotypes |
| Pint | loicpauleve.name/pint | Formal prediction of mutants |
| NuSMV | nusmv.fbk.eu | Formal verification of phenotypes reachability and stability |

### 1.1 This is an executable and reproducible paper

This protocol has been actually edited entirely as a Jupyter notebook before being converted to a LaTeX document for journal-specific editing purposes. The original notebook file is provided as supplemental material. It can also be visualised and downloaded for execution in the CoLoMoTo Interactive Notebook at https://nbviewer.jupyter.org/github/colomoto/colomoto-docker/blob/2018-03-31/usecases/Usecase - Mutations enabling tumour invasion.ipynb.

The blocks beginning with `In [..]` correspond to the content of Jupyter *code cells* which contain the Python instructions to execute. When relevant, the blocks beginning with `Out [..]` display the return value of the last instruction of the related code cell.

Provided Docker and Python are installed, the CoLoMoTo Interactive notebook can be installed by typing and executing the following command^4^ on GNU/Linux, macOS, and Microsoft Windows:

~~~
pip install -U colomoto-docker
~~~

Once installed, the notebook can be executed by typing

~~~
colomoto-docker -V 2018-03-31
~~~

The execution of this command will open a web page with the Jupyter notebook interface, enabling the loading and execution of the code. Note that ”SHIFT+ENTER” must be used to execute each code cell. More information on `colomoto-docker` usage can be obtained by typing `colomoto-docker --help` and by visiting https://github.com/colomoto/colomoto-docker.

### 1.2 Notebook preparation

This notebook makes use of the following Python modules:

~~~
In [1]: import ginsim
          import biolqm
          import maboss
          import pypint
          from colomoto_jupyter import tabulate *# for fixpoint table display*
          from itertools import combinations *# for iterating over sets*
          import matplotlib.pyplot as plt *# for modifying plots*
~~~

## 2 Model

We analyse a Boolean network model of the switch between apoptosis and cell tumour invasion from Cohen et al. [4]. This model can be loaded directly from the GINsim model repository at http://ginsim.org/models_repository.

We first show how to use GINsim [10] to fetch and parse the GINML file (GINsim graph-based XML format, encapsulated in a zginml archive) and display the regulatory graph of the network. To load the model, we copied the URL of the .`zginml` file from the model repository page at http://ginsim.org/node/191.

~~~
In [2]: lrg = ginsim.load("http://ginsim.org/sites/default/files/SuppMat_Model_Master_Model.zginml")
~~~

The regulatory graph (using the graphical setting specified in the model file) can be displayed with the following command:

~~~
In [3]: ginsim.show(lrg)
~~~

The resulting graphics is reproduced in Figure 1.

**Figure 1:**
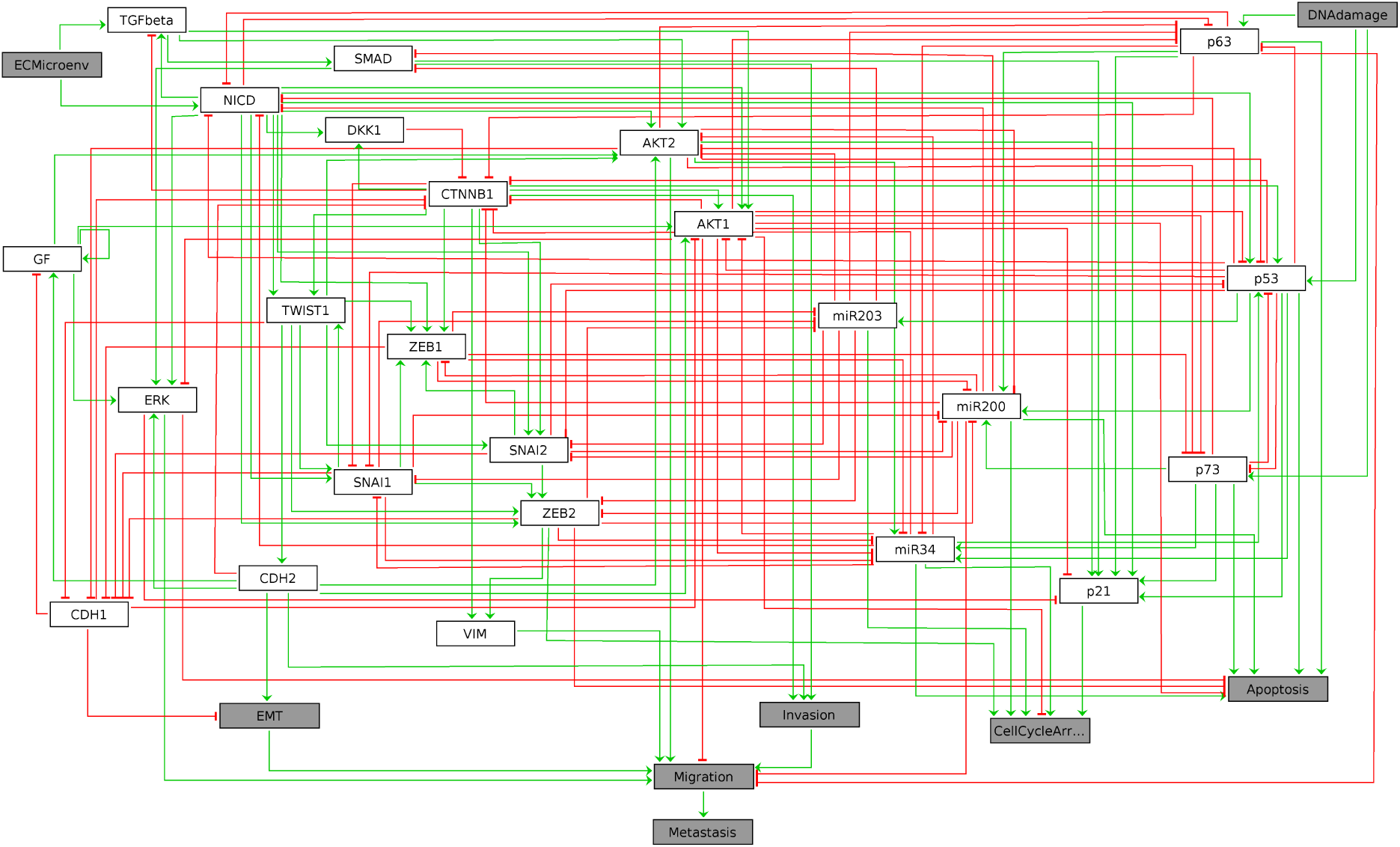
Graphical output resulting from the input code: In [3]: ginsim.show(lrg)

In this regulatory graph, the grey boxes denote input and output vertices (nodes). Green arrows and red T arrows respectively denote activatory and inhibitory interactions. A set of rules combining the vertices with the Boolean operators NOT, AND, and OR, which must be consistent with the regulatory graph, then allows the computation of enabled transitions for each network state. These rules have been defined in Cohen et al. [4] and are specified within the GINsim model.

## 3 Identification of stable states

First, we compute the complete list of logical stable states (or fixpoints) of the model using the java library `bioLQM` [8]. We thus need to convert the GINsim model into bioLQM:

~~~
In [4]: lqm = ginsim.to_biolqm(lrg)
~~~

At that stage, `lrg` is a Python object representing the model suitable for GINsim, and `lqm` is a Python object representing the equivalent model suitable for bioLQM.

The list of stable states of a bioLQM model is computed as follows:

~~~
In [5]: fixpoints = biolqm.fixpoints(lqm)
~~~

Here, `fixpoints` is a Python list of states. A state is encoded as a Python association table (dictionary) which maps each node of the network to a value.

For a nice display of the list of stable states, one can use the `tabulate` function provided in the `colomoto_jupyter` Python library, imported at the beginning of the notebook:

~~~
In [6]: tabulate(fixpoints)
~~~

Figure 2 shows the table as displayed in the notebook. The complete table is given in supplemental data.

**Figure 2:**
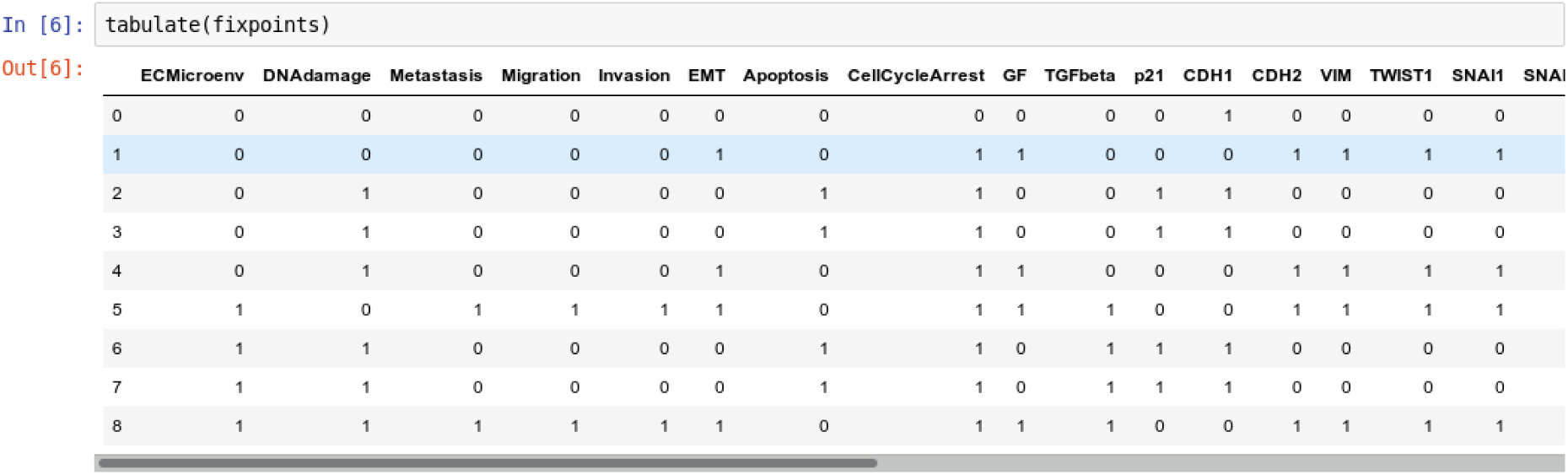
Graphical output resulting from the input code: In [6]: tabulate(fixpoints)

It results that the model has nine stable states, each corresponding to a row in the table, four of which enabling apoptosis (rows with value 1 in fourth column ”Apoptosis”). Note that the input node `DNAdamage` is also active in each of these four states.

A state can be visualised on the regulatory graph using GINsim. For example, the third stable state can be displayed using the following command:

~~~
In [7]: ginsim.show(lrg, fixpoints[2])
~~~

The resulting graphics is reproduced in Figure 3.

**Figure 3:**
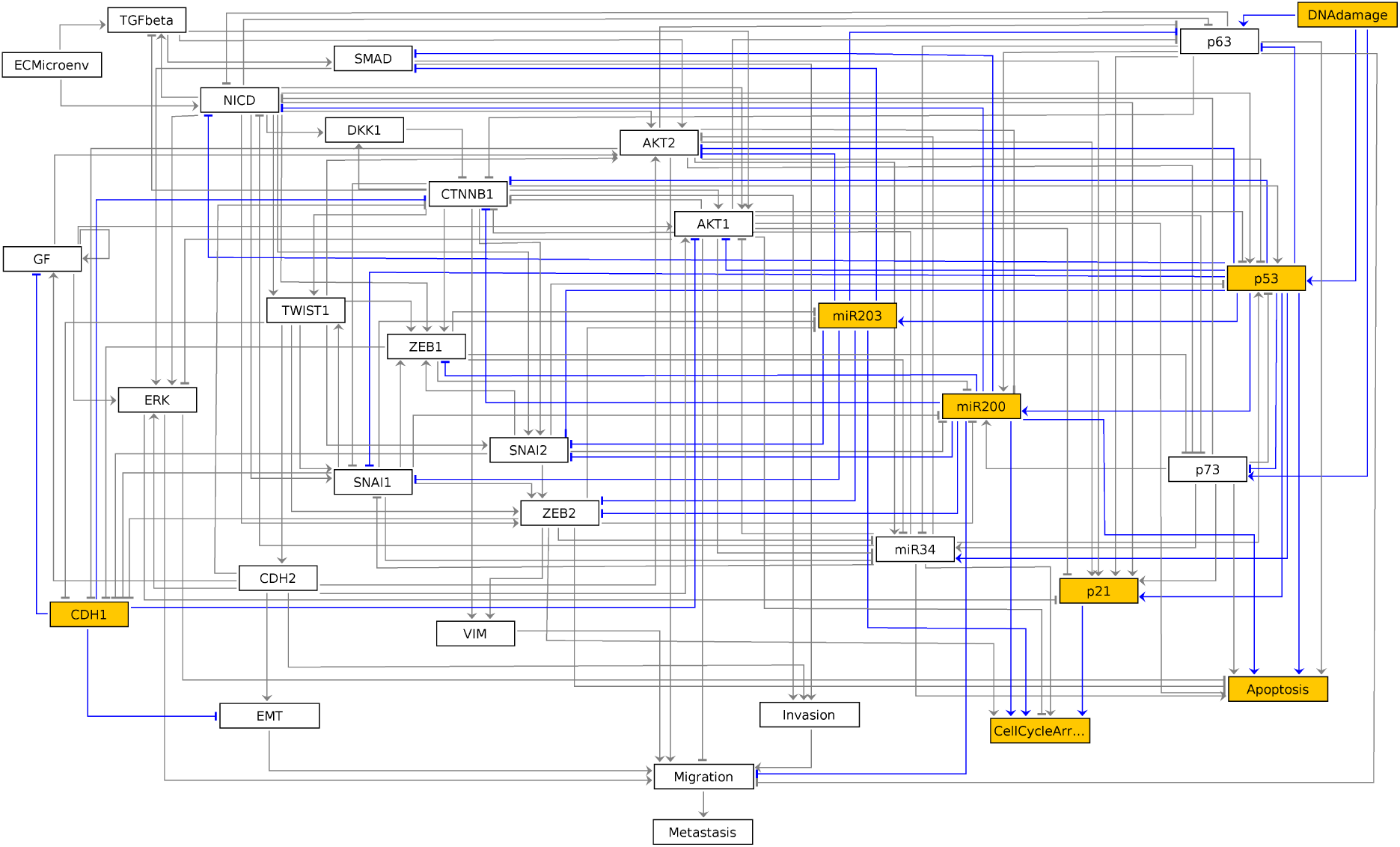
Graphical output resulting from the input code: In [7]: ginsim.show(lrg, fixpoints[2])

In this graph, the vertices shown in white or orange denote components that are OFF (value 0) or ON (value 1) respectively.

## 4 Assessing the probabilities to reach alternative attractors using MaBoSS

MaBoSS [13] is a C++ software that performs stochastic simulations of a Boolean network by translating it into a continuous time Markov processes. Each node activation and inactivation is associated with an up and a *down rate*, which specify the propensity of the corresponding transition. From a given state, the simulation integrates all the possible node updates and derives from their rate a probability and a duration for each transition. By default, all transitions are assigned the same rate. For a given set of initial conditions, MaBoSS produces time trajectories and estimates probabilities of given model states over the whole simulation time. Steady state distributions can thus be approximated, provided that a sufficient number of sufficiently long simulations have been performed.

The aim of this section is to reproduce part of the results obtained by Cohen et al. [4], which show that a Notch (NICD) gain-of-function together with a p53 loss-of-function prevent reaching a stable apoptotic phenotype.

First, we convert the bioLQM model to MaBoSS:

~~~
In [8]: wt_sim = biolqm.to_maboss(lqm)
~~~

The variable `wt_sim` is a Python object that gathers both the Boolean network rules and the settings for the simulations, including the transition rates.

~~~
In [9] : maboss.wg_set_istate(wt_sim)
~~~

### 4.1 Simulation setup

The stochastic simulation of Boolean networks using MaBoSS requires the specification of several parameters.

#### 4.1.1 Initial states

First, a distribution of initial states must be specified: each simulation then starts from a state sampled from this distribution. The distribution is determined by assigning a probability to start in state `0` or in state `1` to each node. By default, a node has a probability `1` to start in state `0`.

The `maboss` Python library provides *widgets* to ease the assignment of this initial distribution. The following code enables the definition of a distribution of initial states with all nodes at `0`, but `DNAdamage` and `ECMicroenv` with equiprobable `0` and `1` values. After pressing ”OK”, the notebook cell will be replaced by the actual Python call resulting in equal probabilities for these two nodes to start in active or inactive states.

~~~
In [9]: maboss.wg_set_istate(wt_sim)
~~~

The notebook will then display the widgets reproduced in Figure 4. The selection of nodes and of initial conditions shown in this figure are then translated in the following code:

~~~
In [10]: *#maboss.wg_set_istate(wt_sim)*
            maboss.set_nodes_istate(wt_sim, ["DNAdamage", "ECMicroenv"], [0.5, 0.5])
~~~

**Figure 4:**
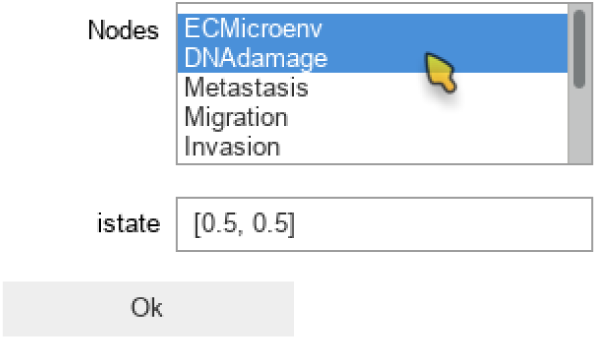
Graphical output resulting from the input code: In [9]: maboss.wg set istate(wt sim)

#### 4.1.2 Output nodes

Using MaBoSS, we can focus on the *output* nodes and ignore the other nodes, which enable us to identify the corresponding phenotypes. This can be done using the following code:

~~~
In [11]: *#maboss.wg_set_output(wt_sim)*
            wt_sim.network.set_output(('Metastasis', 'Migration', 'Invasion',
                                                                        'EMT', 'Apoptosis', 'CellCycleArrest'))
~~~

#### 4.1.3 Simulation parameters

The `update_parameters` method can be used to specify several parameters for the stochastic simulation algorithm. We show below the complete list of parameters with the values obtained by default when translating a model from GINsim. The method can be called with any subset of these parameters.

Among the parameter list, `sample_count` corresponds to the number of simulations performed to compute statistics, while `max_time` is the maximum (simulated) duration of a trajectory. Note that for a proper estimation of probabilities of the stable states, `max_time` needs to be long enough for the simulation to reach an asymptotic solution.

~~~
In [12]: wt_sim.update_parameters(discrete_time=0, use_physrandgen=0,
                      seed_pseudorandom=100, sample_count=50000,
                      max_time=50, time_tick=0.1, thread_count=4,
                      statdist_traj_count=100, statdist_cluster_threshold=0.9)
~~~

### 4.2 Simulation of the wild-type model

The object `wt_sim` represents the input of MaBoSS, encompassing both the network and simulation parameters. The simulations are triggered with the `.run()` method and return a Python object for accessing the results.

~~~
In [13]: %time wt_results = wt_sim.run()
CPU times: user 2.47 ms, sys: 5.02 ms, total: 7.49 ms
Wall time: 6.2 s
~~~

The resulting object gives access to the output data generated by MaBoSS. It includes notably the mean probability over time for the activity of the output states integrated over all the performed simulations.

The function `plot_piechart` displays proportionnaly the mean probability of each output state at the *last* time point. Provided the simulation time has been set high enough, this gives an approximation of the probabilities of the stable states reachable from the specified initial conditions.

~~~
In [14]: wt_results.plot_piechart()
~~~

The resulting graphics is reproduced in Figure 5.

**Figure 5:**
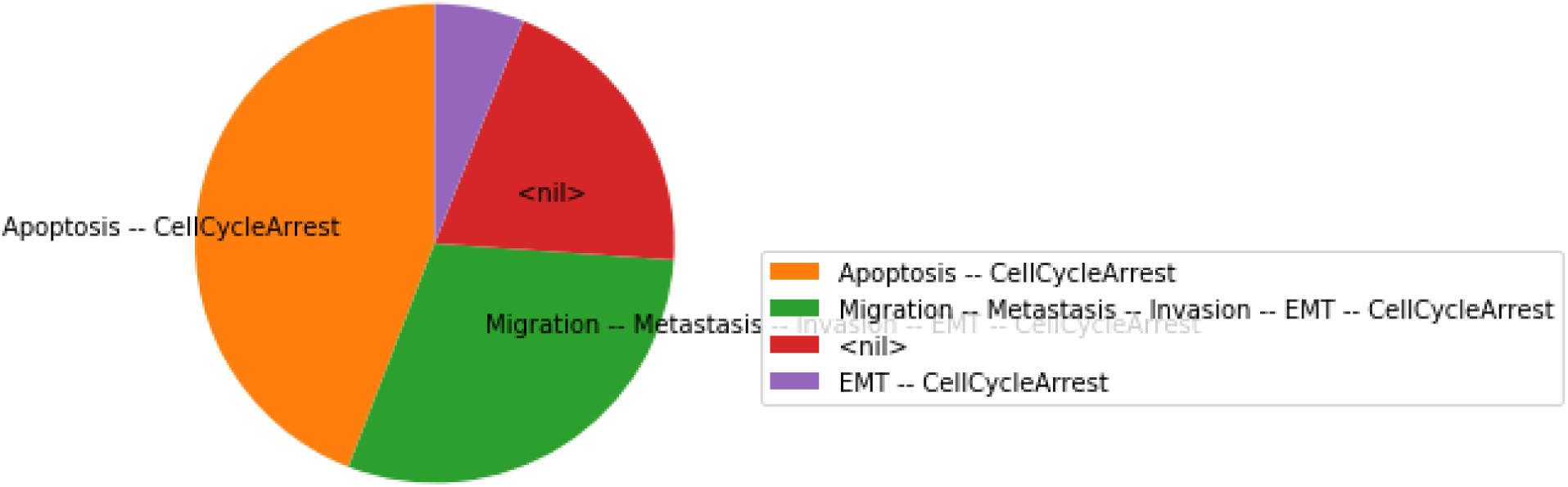
Graphical output resulting from the input code: In [14]: wt results.plot piechart()

In this chart, a state is described by the set of its active output nodes and is associated to a phenotype. For instance, the ”<`nil`>” phenotype has all output nodes set to 0, which was referred to as the ‘’homeostatic state” in the original article; in the case of the ”`Apoptosis — CellCycleArrest`” phenotype, the two output nodes `Apoptosis` and `CellCycleArrest` are simultaneously active, while the other output nodes are inactive; the ” `EMT -- CellCycleArrest`” phenotype denotes cells that have gone through the epithelial to mesenchymal transition, but did not invade the tissue, hence the output nodes `Invasion`, `Migration` and `Metastasis` are inactive; finally the ”`Migration -- Metastasis -- Invasion -- EMT -- CellCycleArrest`” phenotype corresponds to a metastatic state, *i.e*. to cells that went through EMT, invaded the tissue and migrated to a distant site.

From this plot, we can deduce that, from the specified set of initial conditions, the apoptotic state (orange section), the EMT (purple section) and the metastatic states (green section) can be reached (the proportion of simulations that reached none of these phenotypes correspond to the red section).

The mean value of each output node during the simulations can be plotted with the following command:

~~~
In [15]: wt_results.plot_node_trajectory()
~~~

The resulting graphics is reproduced in Figure 6.

**Figure 6:**
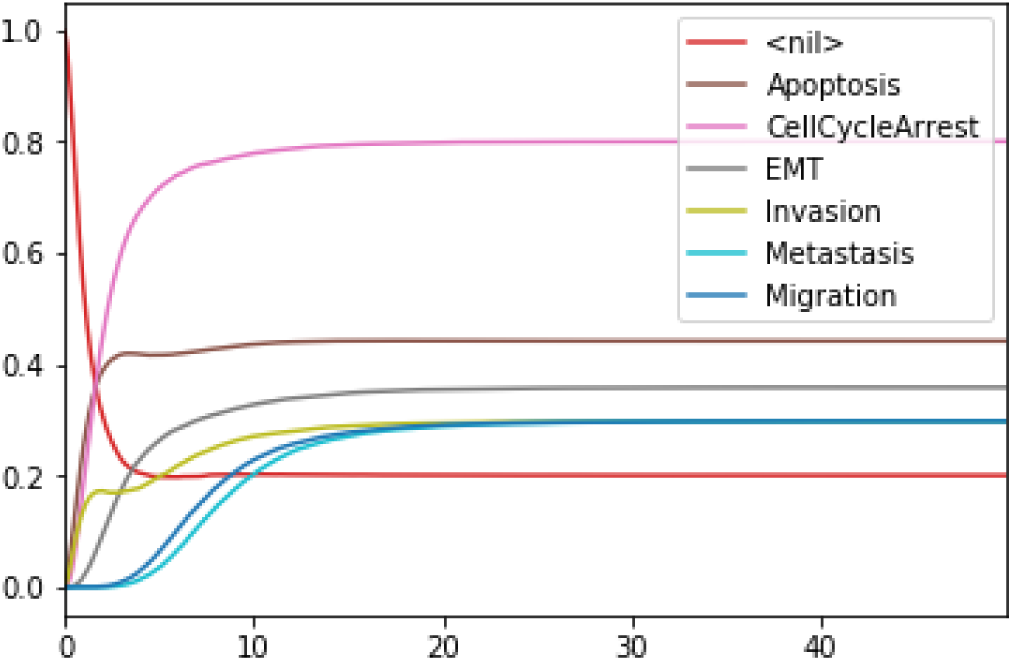
Graphical output resulting from the input code: `In [15]: wt_results.plot_node_trajectory()`

### 4.3 Simulation of double mutant Notch++/p53--

In the original article [4], the authors analysed the double Notch++/p53-- mutant, *i.e.*, the combination of a Notch gain-of-function combined with a p53 loss-of-function, showing that all trajectories lead to a metastatic state.

A mutant can be configured by copying the wild-type model, and use the `mutate` method to model the desired gains and losses of function:

~~~
In [16]: mut_sim = wt_sim.copy()
            mut_sim.mutate("p53", "OFF")
            mut_sim.mutate("NICD", "ON")
~~~

The modified model can then be simulated exactly as for the wild-type case:

~~~
In [17]: %time mut_results = mut_sim.run()
CPU times: user 4.22 ms, sys: 4.98 ms, total: 9.2 ms
~~~

~~~
Wall time: 5.98 s
~~~

~~~
In [18]: mut_results.plot_piechart()
~~~

The resulting graphics is reproduced in Figure 7.

**Figure 7:**
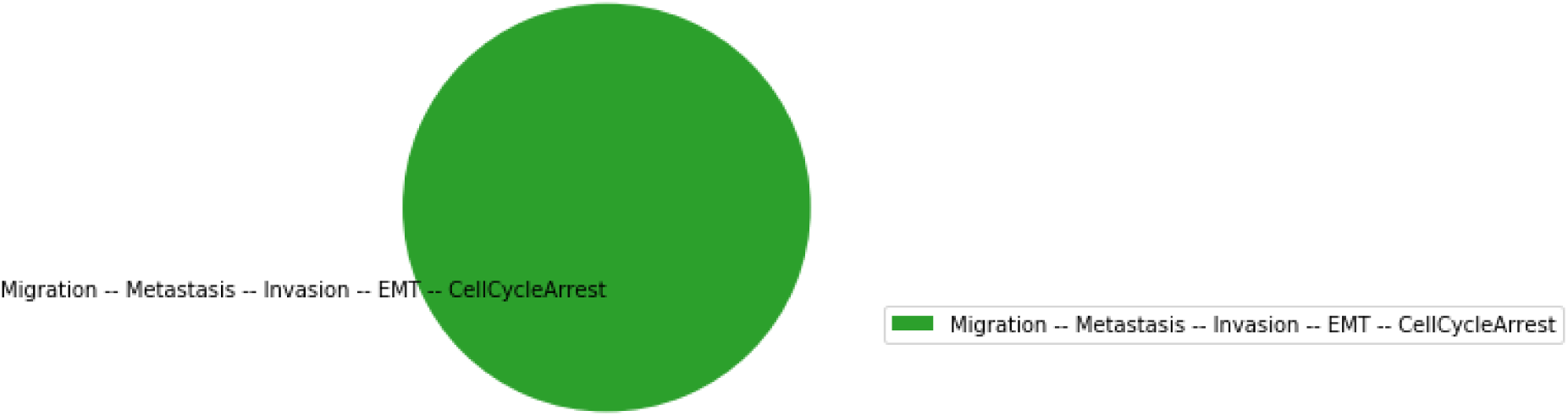
Graphical output resulting from the input code: `In [18]: mut results.plot piechart()`

Hence, using the same parameters as for the wild-type model, all the trajectories obtained for the double mutant model reach the metastatic invasive state exclusively. This suggests that such a double mutation can be responsible for a loss of apoptotic capability of cancer cells.

## 5 Formal analysis with Pint and NuSMV

In the above section, the conclusion regarding the loss of apoptotic stable state relies on stochastic simulations, which, in general, may not offer a complete coverage of the possible trajectories. Therefore, one may want to formally verify whether the loss of reachable stable apoptosis state is total or not. First, we show how to use Pint [12] to predict combinations of mutations which are guaranteed to prevent the activation of apoptosis. Next, we use the software NuSMV [2] to evaluate formally the Notch++/p53-- double mutant. Finally, we use MaBoSS to assess the efficiency of new combinations of mutations predicted by Pint.

### 5.1 Formal predictions of mutations from the wild-type model

Pint implements formal methods which allows deducing combinations of mutations which are guaranteed to block the reachability of a given state.

First, we convert the bioLQM model to Pint:

~~~
In [19]: an = biolqm.to_pint(lqm)
~~~

Then, we transfer the initial conditions defined in MaBoSS to the `an` Pint model. Like MaBoSS, Pint supports multiple initial values for a single node. However, in contrast to MaBoSS, Pint does not consider probability distributions.

~~~
In [20]: an.initial_state.update(wt_sim.get_initial_state())
            an.initial_state.changes() *# display non-default (0) initial value*
~~~

~~~
Out[20]: {'DNAdamage': (0, 1), 'ECMicroenv': (0, 1)}
~~~

Given a (partial) state specification, Pint provides the method `oneshot_mutations_for_cut`, which returns different sets of mutations guaranteed to prevent any trajectory from any possible initial state to reach, *even transiently*, the specified state.

~~~
In [21]: %time an.oneshot_mutations_for_cut(Apoptosis=1, \
exclude={"ECMicroenv", "DNAdamage"})
CPU times: user 8.74 ms, sys: 5.04 ms, total: 13.8 ms
Wall time: 298 ms
~~~

~~~
Out[21]: [{'ZEB2': 1},
               {'AKT1': 1},
               {'AKT2': 1},
               {'ERK': 1},
               {'NICD': 1, 'SNAI2': 1, 'ZEB1': 1},
               {'SNAI2': 1, 'ZEB1': 1, 'p63': 0},
               {'SNAI2': 1, 'ZEB1': 1, 'miR203': 1},
               {'NICD': 1, 'SNAI2': 1, 'p73': 0},
               {'SNAI2': 1, 'p63': 0, 'p73': 0},
               {'SNAI2': 1, 'miR203': 1, 'p73': 0},
               {'NICD': 1, 'ZEB1': 1, 'p53': 0},
               {'ZEB1': 1, 'p53': 0, 'p63': 0},
               {'ZEB1': 1, 'miR203': 1, 'p53': 0},
               {'NICD': 1, 'p53': 0, 'p73': 0},
               {'p53': 0, 'p63': 0, 'p73': 0},
               {'miR203': 1, 'p53': 0, 'p73': 0}]
~~~

Among the returned mutation sets, one can spot the mutation {'`NICD`' `: 1`, '`p53`' `: 0`, '`p73`': `0`}, which combines a gain-of-function of Notch (' NICD': 1) with a loss-of-function of p53 ('`p53`': 0), along with a loss-of-function of p73 ('`p73`': `0`).

Noteworthy, forbidding *transient* reachability entails a stronger constraint than just preventing any *stable* state with the specified property. Indeed, some mutations may remove the stability of the specified states, while some trajectories may still traverse some of them, but only transiently.

Therefore, the sets of mutations returned by Pint, albeit correct, might be non-minimal for controlling only the long-term dynamics of the system. Finally, note that the analysis of Pint can give incomplete results. This is due to the technology on which the computation relies (static analysis), which allows addressing very large scale networks.

### 5.2 Revisiting the Notch++/p53-- double mutant

We will first formally analyse the Notch++/p53-- double mutant to show that asymptotic apoptosis is forbidden, although transient activation of apoptosis node might still be possible.

One can apply a mutation on a Pint model using the `lock` method. A new model is returned with a constant value for the corresponding nodes.

~~~
In [22]: mut_an = an.lock(NICD=1, p53=0)
~~~

Then, we use the temporal logic CTL [3] to specify formally the dynamical properties to verify. CTL expression can be built using the `colomoto.temporal_logics` Python module.

~~~
In [23]: from colomoto.temporal_logics import *
~~~

First, the existence of a trajectory leading to a *transient* state where Apoptosis is active can be specified as follows:

~~~
In [24]: transient_apoptosis = EF(S(Apoptosis=1))
~~~

`EF` is a temporal logic operator that is true if there exists at least one trajectory leading to a state verifying the properties given as argument. Here the property `S(Apoptosis=1)` specifies that the state has the node `Apoptosis` active.

Next, the existence of a trajectory leading to a *stable* `Apoptosis` activation can be specified as follows:

~~~
In [25]: stable_apoptosis = EF(AG(S(Apoptosis=1)))
~~~

Here, `AG` enforces that *all* the states reachable via any trajectory have the node `Apoptosis` active. Finally, we gather these two properties in a Python dictionary for later use:

~~~
In [26]: ctl_specs = {
                  "reach-apoptosis": transient_apoptosis,
                  "stable-apoptosis": stable_apoptosis
}
~~~

The adequation of a model with a CTL property can be assessed using a *model-checker* such as NuSMV [1].

Pint provides a conversion to NuSMV models. By default, the NuSMV model considers any initial state. With the `skip_init=False` option, we enforce that the properties are verified only from the initial states defined earlier.

~~~
In [27]: smv = mut_an.to_nusmv(skip_init=False)
~~~

We then add the properties defined above, and ask NuSMV to verify them.

~~~
In [28]: smv.add_ctls(ctl_specs)
            %time smv.verify()
CPU times: user 2.82 ms, sys: 6.02 ms, total: 8.84 ms
Wall time: 20 s
Out[28]: {'reach-apoptosis': True, 'stable-apoptosis': False}
~~~

Interestingly, the Notch++/p53-- double mutant can still reach an apoptotic state, but only transiently: the property `stable-apoptosis` being false, it is guaranteed that all trajectories eventually lead to stable apoptosis inactivation.

To complete our analysis, we now consider the triple mutant obtained by adding a loss-of-function of `p73`. As predicted by Pint, transient reachability of apoptosis is impossible in this triple mutant. We can use NuSMV to further verify that it is the case, using the following code:

~~~
In [29]: smv_mut3 = an.lock(NICD=1, p53=0, p73=0).to_nusmv(skip_init=False)
            smv_mut3.add_ctls(ctl_specs)
            smv_mut3.verify()
Out[29]: {'reach-apoptosis': False, 'stable-apoptosis': False}
~~~

### 5.3 Analysis of formally predicted SNAI2++/ZEB1++/miR203++ triple mutant

The mutant combinations predicted with Pint should be refined when the aim is to control specifically stable behaviours. In general, given a set of mutations guaranteed to block any transient activation of a node, one may verify whether only a subset of them are sufficient to achieve proper control of the sole stable states.

We show here how we can take advantage of the Python environment to provide a small program, which, for each subset of mutations of a multiple mutant (here a triple gain-of-function for SNAI2, ZEB1 and miR203), performs stochastic simulations with MaBoSS to assess the probabilities to reach the different stable behaviours from the specified set of states.

The computation can take a couple of minutes. The results are shown in a graphical form (coloured pie charts) for each single and double loss-of-function combination.

~~~
In [30]: formal_mutant = {'SNAI2': 1, 'ZEB1': 1, 'miR203': 1}
             for i in [1, 2]:
                   *# for any subset of mutations of size 1 then 2*
                   for mutants in combinations(formal_mutant, i):
                        # *copy the wild-type MaBoSS model*
                        masim = wt_sim.copy()
                        # *apply the mutations*
                        for m in mutants:
                             masim.mutate(m, "ON" if formal_mutant[m] else "OFF")
                        # *run the simulations*
                        mares = masim.run()
                        # *plot the piechart of stable states*
                        mares.plot_piechart(embed_labels=False, autopct=4)
                        # *print the mutation in the title*
                        def mutname(m):
                              return m + ("++" if formal_mutant[m] else "—")
                        name = "/".join(map(mutname, mutants))
                        plt.title("%s mutant" % name)
~~~

The resulting graphics is reproduced in Figures 8 to 13.

**Figure 8:**
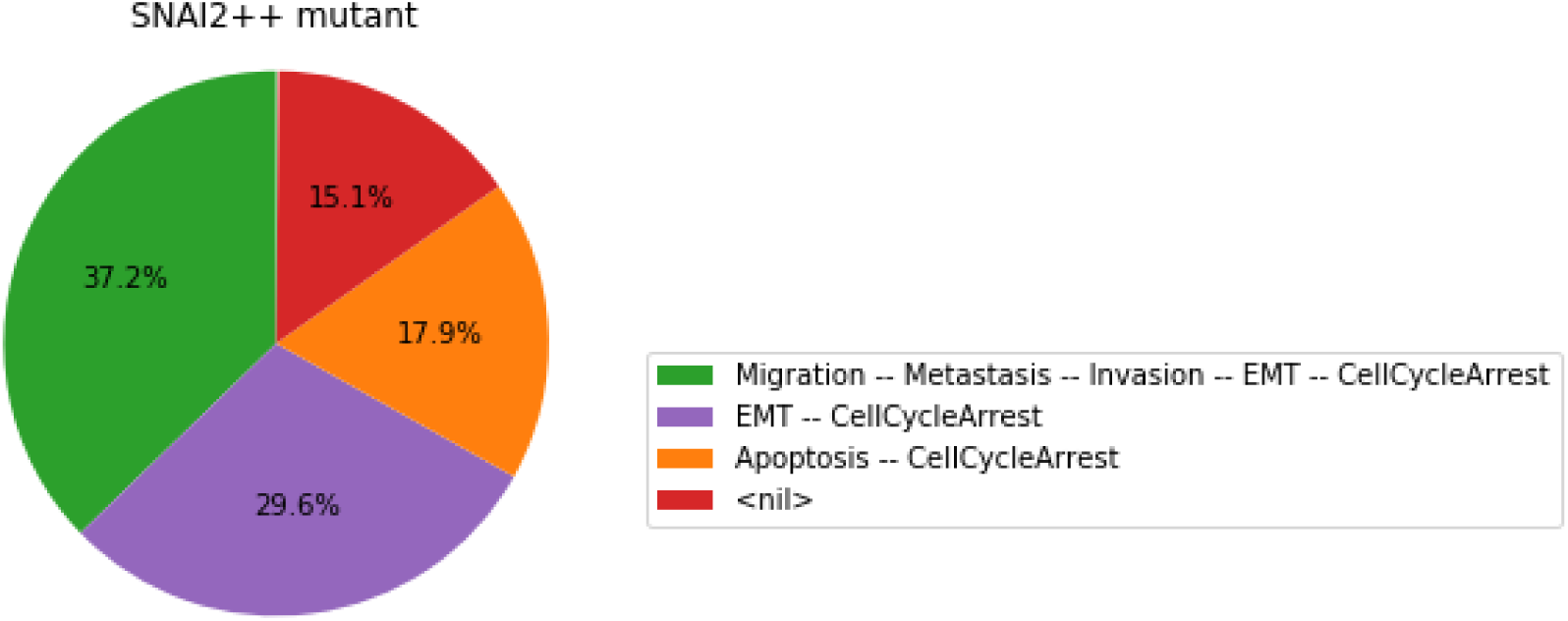
Graphical output resulting from the input code: `In [30]`

**Figure 9:**
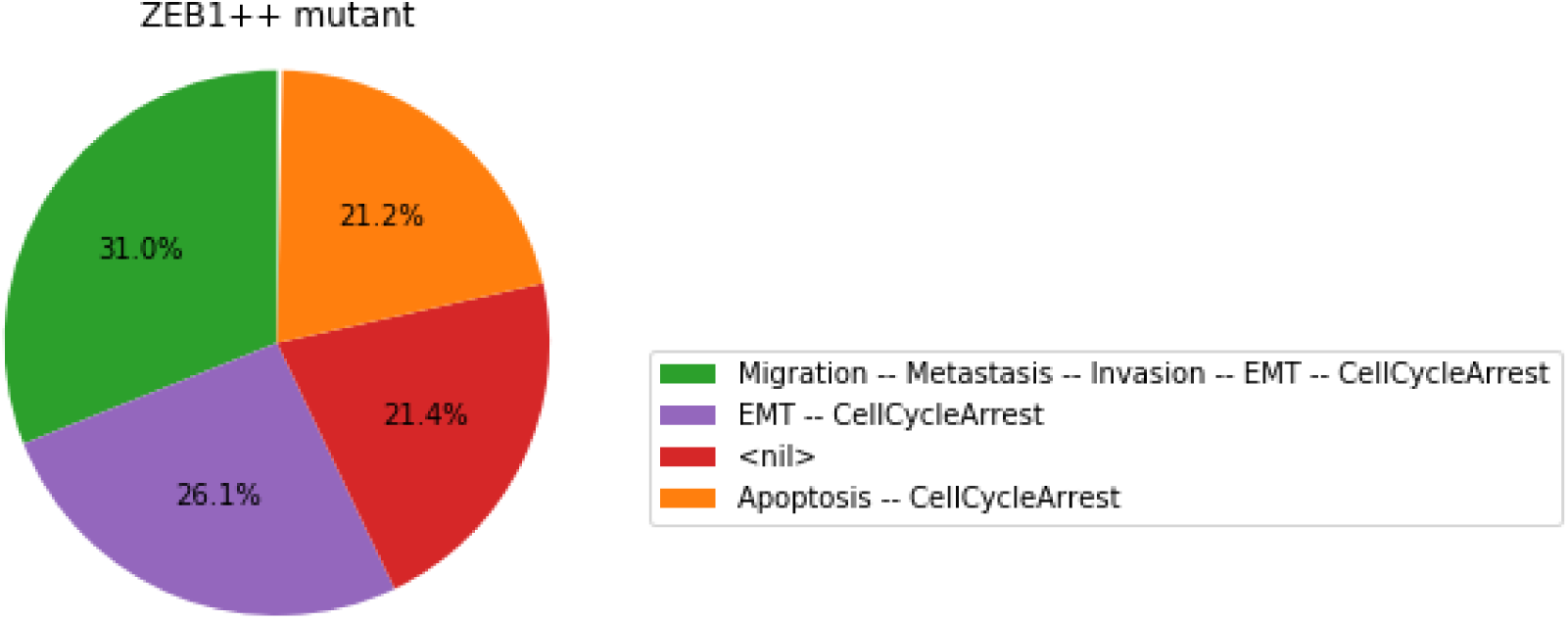
Graphical output resulting from the input code: `In [30]`

**Figure 10:**
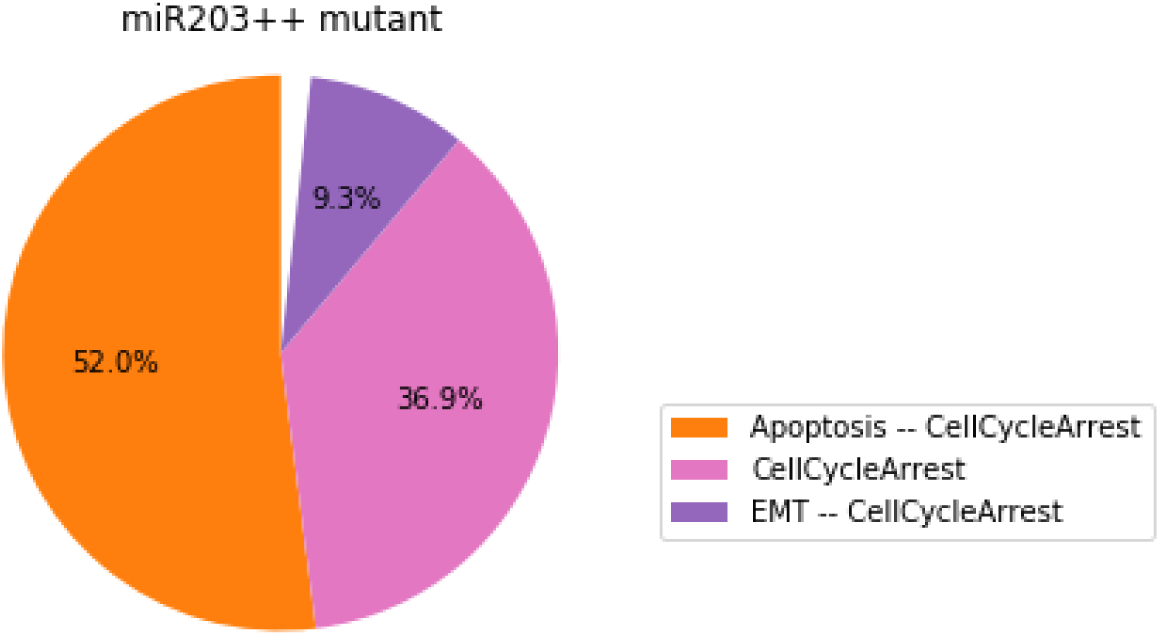
Graphical output resulting from the input code: `In [30]`

**Figure 11:**
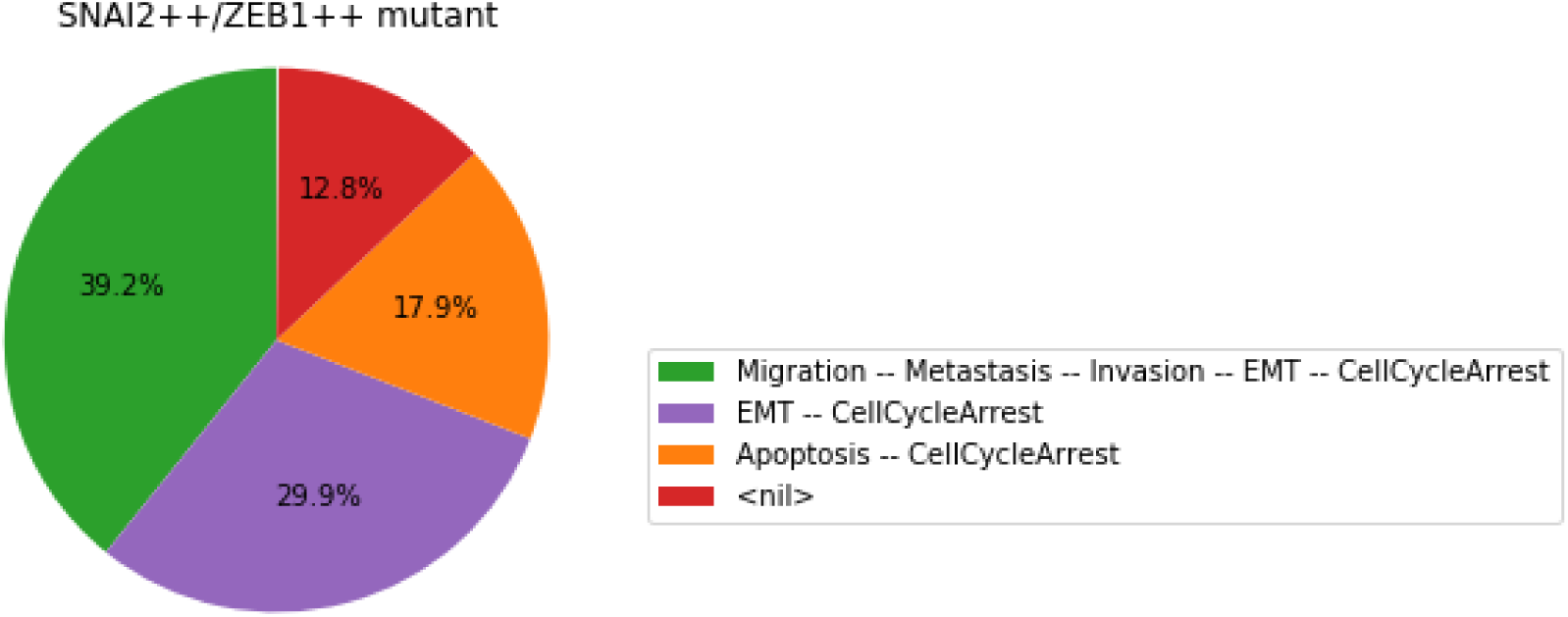
Graphical output resulting from the input code: `In [30]`

**Figure 12:**
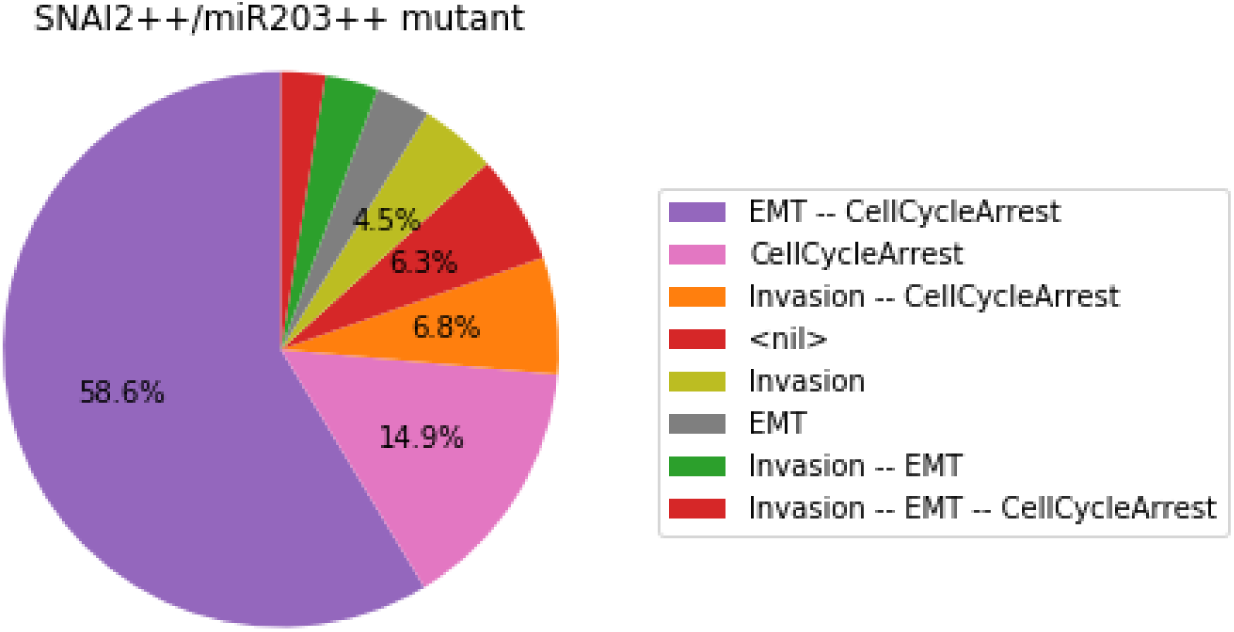
Graphical output resulting from the input code: `In [30]`

**Figure 13:**
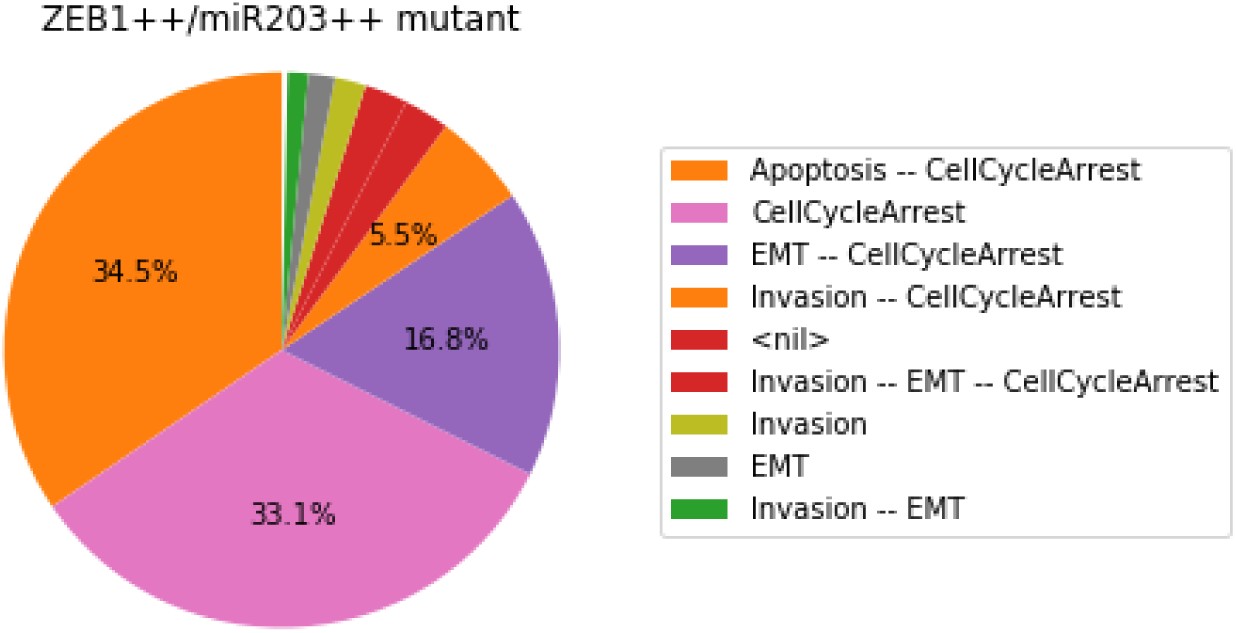
Graphical output resulting from the input code: `In [30]`

Note that only one of the pie charts shows an absence of apoptotic state: the SNAI2++/miR203++ double mutant (Figure 12).

This can be formally verified with NuSMV, as we did for the Notch++/p53-- mutant:

~~~
In [31]: smv_mut_test = an.lock(SNAI2=1, miR203=1).to_nusmv(skip_init=False)
            smv_mut_test.add_ctls(ctl_specs)
            smv_mut_test.verify()
Out[31]: {'reach-apoptosis': True, 'stable-apoptosis': False}
~~~

## 6 Conclusion

With this notebook, we showed how the Python interface and Jupyter integration of GINsim, bioLQM, MaBoSS, and Pint ease the delineation of sophisticate re-executable computational analyses of qualitative models of biological networks, combining and chaining different software with a unified interface.

In this protocol, we demonstrated the assets of this framework by revisiting the analysis of a Boolean model of the cell fate decision between apoptotic and metastatic phenotypes, initially defined with GINsim. We could thereby reproduce results previously obtained with GINsim and MaBoSS, which demonstrate that the Notch++/p53-- double mutant can suppress the apoptotic outcome. Furthermore, our formal analysis of trajectories using Pint enabled us to deduce novel ”anti-apoptotic” combinations of mutations, including a triple mutant that forbids even transient activation of apoptosis, which have been subsequently quantified using MaBoSS.

The resulting combinations of mutations point to potential synergistic genetic interactions underlying uncontrolled tumour proliferation. These combinations would deserve further analysis, in particular regarding potential correlations with specific clinical outcomes. For example, one could check whether the loss of apoptosis triggering correlates with higher tumour grades.

Similar computational analyses could be performed to predict combinations of perturbations enforcing the existence of a given stable phenotype, *e.g*. apoptosis, which could then serve as a basis to design novel therapeutic strategies.

## Author Contributions

NL, AN, CH, LP implemented the necessary Python modules, their integration in the Jupyter interface, and the Docker image. NL, AN, GS, DT, AZ, LC, LP participated to the general design of the tutorial notebook. All authors participated to the writing of the article.

## Footnotes

1 Available at https://github.com/colomoto/colomoto-docker

2 https://docker.com

3 https://jupyter.org

4 You may have to use `pip3` instead of `pip` depending on your configuration

